# Sub-sarcomeric regulation of thin and thick filaments in skeletal muscle myofibrils

**DOI:** 10.1101/2025.11.21.689690

**Authors:** Kristina Sliogeryte, Luca Fusi

## Abstract

Muscle contraction relies on the coordinated activation of myosin motors from a folded-OFF state on the thick filament surface and their actin tracks on the thin filaments in response to calcium. Thick filaments contain distinct regulatory zones defined by the presence of myosin-binding protein C (MyBP-C) and titin super-repeats, but the control of myosin OFF/ON states within these zones has not been directly resolved. Here we do so, by fluorescence polarization microscopy (FPM). Using orientation-specific probes on myosin we show that folded-OFF motors are enriched in the MyBP-C–containing C zone in relaxed myofibrils. Under titin-based passive tension or partial calcium activation, active motors are enriched in the D zone at the filament tips, which lacks MyBP-C. Troponin probes further reveal that myosin enhances thin-filament activation in the region of filament overlap and drives activation into adjacent non-overlap regions. These findings uncover zone-specific control of myofilament activation within the sarcomere and establish FPM as a powerful tool for investigating disease-linked myofilament protein variants and therapeutic modulation.

## Introduction

Muscle contraction requires the coordinated activation of actin-containing thin filaments and myosin-containing thick filaments within the sarcomere, the fundamental contractile unit of striated muscle, initiated by calcium release in the cytoplasm (1–3). In resting muscle, when calcium is absent, the thin filament is switched off by troponin and tropomyosin, which block the myosin-binding sites on actin (4–6). Concomitantly, most myosin motors on the thick filament are unavailable for contraction because they fold back onto the filament surface in the interacting-heads motif (IHM) (7, 8). This structural configuration is typically associated with a biochemical state of myosin characterized by markedly reduced ATP turnover, known as the super-relaxed (SRX) state (9).

Upon activation, calcium binding to troponin partially shifts tropomyosin on the thin filament, exposing myosin-binding sites and allowing constitutively ON myosin motors to bind to actin in the region of filament overlap (A-band; Fig. 1A), thereby generating force (10, 11). The resulting stress activates a mechano-sensing feedback mechanism that promotes the rapid transition of folded-OFF myosin motors into the ON state (10), progressively increasing the population of force-generating motors. This, in turn, further enhances thin filament activation within the A-band (12), although how myosin influences thin filament regulation in regions lacking filament overlap (I-band; Fig. 1A) remains unclear.

**Figure 1.**
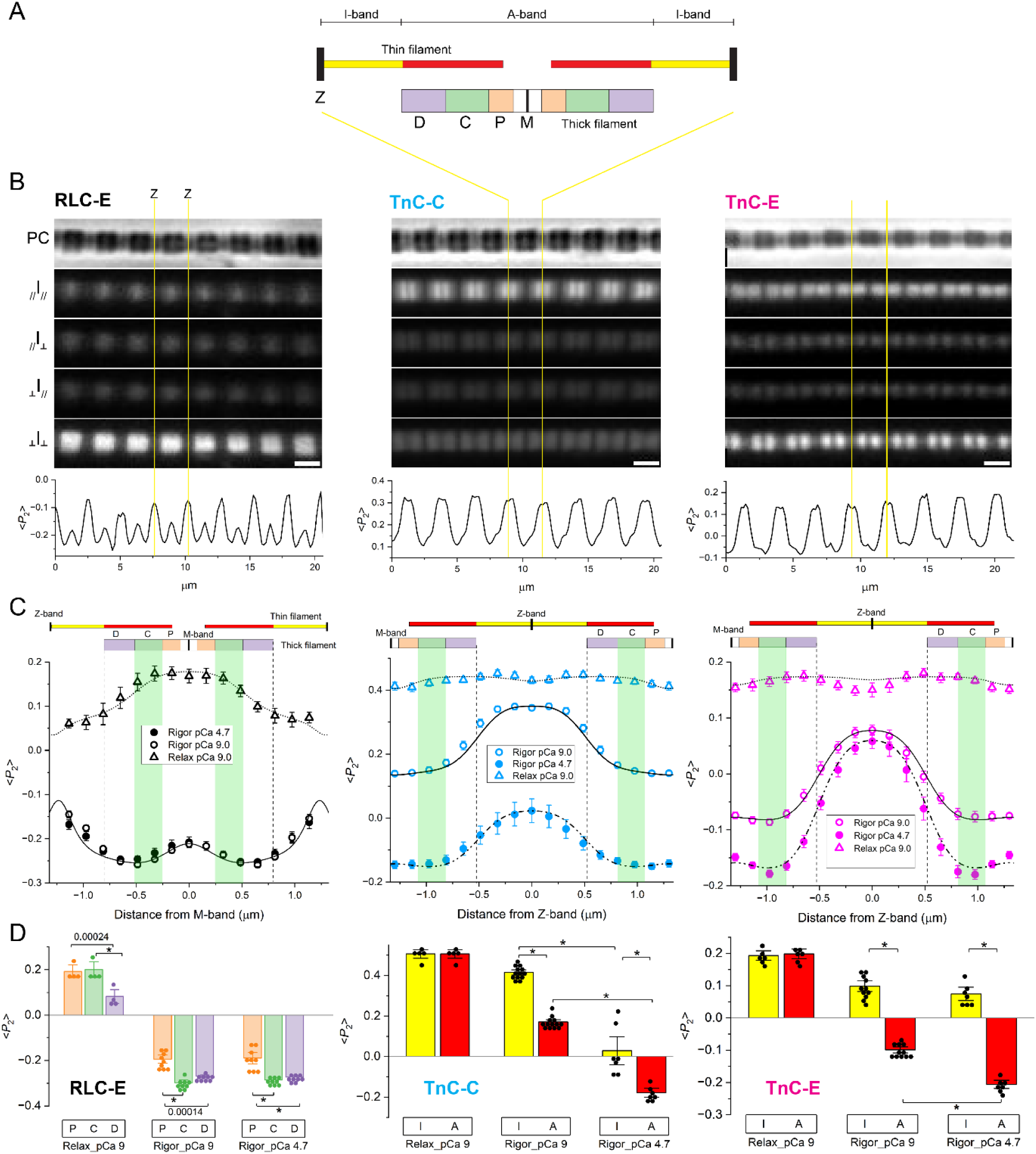
Spatial distributions of TnC and RLC orientations in thin- and thick-filament zones in rigor and relaxed myofibrils. A) Schematic of a sarcomere highlighting thin (I,A) and thick (P, C, D) filament zones. Sarcomere ends, Z-band (Z); sarcomere midpoint, M-band (M). B) Phase contrast (PC) and Fluorescence Polarization Microscopy (FPM) images of myofibrils exchanged with RLC-E, TnC-C and TnC-E probes in rigor conditions in the absence of calcium (pCa 9). Yellow lines mark the sarcomere ends (Z-bands). Scale bar, 2 µm. One-dimensional distributions of <*P*_2_> along the myofibril axis are shown below the polarised images. C) Average distribution of <*P*_2_> (mean±SE) for the RLC-E probe (black) along the thick filament and for the TnC-C (cyan) and TnC-E (magenta) probes along the thin filament in single sarcomeres, in Rigor-pCa 9 (empty circles), Rigor-pCa 4.7 (filled circles) and in Relaxing conditions at pCa 9 (triangles) at 17°C. RLC-E data: Rigor-pCa 9, sarcomere length (SL)=2.47±0.02 µm (n=9 myofibrils); Rigor-pCa 4.7, SL=2.59±0.03 µm (n=10); Relax-pCa 9, SL=2.58 ± 0.03 µm (n=4). TnC-C data: Rigor-pCa 9, SL=2.63±0.01 µm (n=14); Rigor-pCa 4.7, SL=2.67±0.02 µm (n=7); Relax-pCa 9, SL=2.53±0.06 µm (n=5). TnC-E data: Rigor-pCa 9, SL=2.61±0.03 µm (n=11); Rigor-pCa 4.7, SL=2.65±0.03 µm (n=7); Relax-pCa 9, SL=2.49±0.03 µm (n=6). Vertical dashed lines, thick filament tips; green bands, C-zone. Model simulations of probe orientation are shown as dotted (Relax), solid (Rigor pCa 9) and dashed-dotted (Rigor pCa 4.7) lines. A diagram of the sarcomere at a length of 2.6 µm is shown above each graph. D) Average <*P*_2_> for RLC and TnC probes in thin and thick filament zones determined from the model simulations in C (asterisks indicate P<0.0001). Model parameters are shown in Table S2.

Each half thick filament can be divided into three structurally and functionally distinct domains (Fig. 1A): a central C-zone containing myosin binding protein-C (MyBP-C), flanked by P- and D-zones incorporating distinct repeats of the titin molecule. Variants in MyBP-C and titin are associated with skeletal and cardiac myopathies (13–16), yet how these proteins regulate the OFF-to-ON transition of myosin across filament domains remains poorly understood. Moreover, filament stress during contraction is non-uniform, increasing linearly from the filament tips toward the M-band, which may influence the mechano-sensing transition of myosin within each filament domain (17).

Recent *in vitro* and *in situ* cryo-EM reconstructions of cardiac thick filaments have provided detailed insights into the organization of myosin, titin, and MyBP-C within the C-zone (18–20). These reconstructions show that the C-terminus of MyBP-C stabilizes the folded OFF conformation of myosin through direct interactions with the motors, while its N-terminus forms links with the overlapping thin filament (19). The stabilisation of the myosin OFF state by MyBP-C is consistent with ATP-turnover studies showing enrichment of the SRX myosin population within the C-zone (21, 22), although the equivalence of the structural OFF and biochemical SRX states has been recently questioned (23–26).

While the structure of the D-zone remains unresolved, X-ray interference studies in cardiac muscle have proposed distinct myosin conformations corresponding to different regulatory regions of the thick filament -namely, folded motors arranged in helical tracks within the C-zone and folded, non-helically ordered motors within the D-zone- and have further suggested a sequential activation of myosin motors progressing from D to C to P during contraction (27). However, existing fluorescence- and X-ray-based structural methods are limited, because they provide only ensemble-averaged measurements of the regulatory states of the myofilaments, masking zone-specific differences in filament regulation.

To overcome these limitations, we developed a Fluorescence Polarization Microscopy (FPM) approach that enables direct visualization of sub-sarcomeric structural changes in both thin and thick filaments within individual skeletal muscle myofibrils. Using orientation reporters on the myosin regulatory light chain (RLC), we show that relaxed myofibrils exhibit an enrichment of folded myosin motors in the C-zone, providing direct evidence in situ that MyBP-C stabilizes the myosin OFF state. By resolving zone-specific OFF-to-ON transitions of myosin motors triggered by titin-based passive tension and calcium activation, we show that motors in the P- and C-zones exhibit greater OFF-state stability than those in the D-zone under filament stress. Furthermore, using troponin C (TnC) probes, we reveal the spatial regulation of thin filament activation within the sarcomere, showing that during contraction myosin-dependent structural changes propagate from the A-band to the I-band, likely through a force-dependent mechanism. Together, these findings uncover the spatially coordinated regulation of thin- and thick-filament activation within the sarcomere, which underpins the control of force generation in skeletal muscle. Moreover, our study establishes FPM as a powerful tool for investigating disease-associated alterations in myofilament structure and function at sub-sarcomeric resolution.

## Results

### Mapping the regulatory state of thin and thick filaments in relaxed myofibrils

We mapped the conformation of myosin and troponin along thick and thin filaments, respectively, using FPM on skeletal muscle myofibrils exchanged with bifunctional rhodamine probes on RLC and TnC. The RLC-E probe on the C-lobe of the myosin RLC was used to monitor the ON/OFF conformation of the myosin motors on the thick filament (12, 28), while the TnC-C probe in the N-lobe of TnC and the TnC-E probe on the C-lobe of TnC reported the orientation of the regulatory head and IT-arm of troponin, respectively, in the thin filament (12, 29, 30).

First, we determined the orientation distributions of RLC and TnC probes along thick and thin filaments, respectively, in relaxed myofibrils at 17°C (pCa=9), at ∼400nm spatial resolution. One-dimensional distributions of intensity (*I*) and probe orientation along the myofilament axis, quantified by the order parameter <*P*_2_> (ranging between −0.5 and +1 for probe orientations perpendicular and parallel to the filament axis, respectively (31)), were calculated from the polarised images of the myofibrils (Fig. 1B,C; Fig. S1). The spatial average of <*P*_2_> for the RLC-E probe across a single sarcomere of a myofibril was similar to bulk <*P*_2_> values measured previously in muscle fibres using wide-field Fluorescence for In-Situ Structure (FISS) (28) (Table S1), indicating that the average myosin motor conformation on the thick filament in relaxed myofibrils is similar to that observed in relaxed skeletal muscle fibres. However, FPM revealed a non-uniform spatial distribution of RLC orientations along the thick filament: <*P*_2_> was higher within ±300 nm of the thick filament midpoint (M-band), including the P- and C-zones, and decreased in the D-zone (Fig. 1C, black triangles), indicating that myosin motors in the central regions of the filament adopt orientations more parallel to the filament axis.

The fluorescence intensity of the TnC probes was higher in the I-band (Fig. S1), indicating a more efficient TnC exchange in this filament domain under the rigor conditions used in our exchange protocol (see Methods), consistent with the non-uniform exchange of troponin in rigor myofibrils observed previously (32). Nevertheless, the orientation parameter <*P*_2_> of the TnC probes was unaffected by changes in fluorescence intensity and was uniform along the thin filaments (Fig. 1C, cyan and magenta triangles), indicating a homogeneous troponin conformation in relaxed myofibrils, consistent with bulk <*P*_2_> values obtained by FISS in relaxed muscle fibres (33) (Table S1).

Accurate estimates of the spatial distribution of the orientation of the RLC probe in the P-, C-, and D-zones of the thick filament, as well as the TnC probes in the I- and A-zones of the thin filament, are constrained by the point spread function (PSF) of the microscope. Indeed, the PSF broadens the fluorescence signals beyond the true boundaries of each zone and produces apparent signals in sarcomeric regions where probes are not expected to localize (Fig. 1C; Fig. S1). To address this, zonal values of <*P*_2_> for each probe were derived from model simulations of the orientation distributions, which incorporated the effects of the experimental PSF and a small fraction (10%) of non-specific probe binding (Supplementary Text and Fig.S2). The orientation distributions for TnC-C and TnC-E probes in relaxed myofibrils were simulated with similar values of <*P*_2_> in the A- and I-bands (Fig. 1D; Table S2), indicating a uniform conformation of troponin along the thin filaments. In contrast, the RLC probe orientation distribution was best fit with <*P*_2_> values in the P- and C-zones higher than that in the D-zone (Fig. 1D; Table S2), supporting the conclusion that the fraction of folded OFF motors parallel to the filament axis is higher in P- and C-zones of the thick filament in relaxed myofibrils.

### Thin Filaments exhibit distinct regulatory states in the I- and A-bands of the sarcomere in rigor myofibrils

Next, we studied the effect of myosin binding to actin on the regulatory state of the thin filament in rigor myofibrils under zero-force conditions. In the absence of calcium (pCa 9), the <*P*_2_> values for the RLC-E probe became more negative across all zones of the thick filament (Fig. 1C, black open circles), indicating a more perpendicular orientation of the actin-attached myosin motors relative to the filament axis. The average value of <*P*_2_> was similar to that measured by FISS in muscle fibres in rigor (Table S1). However, FPM analysis showed that <*P*_2_> for the RLC-E probe in the P-zone was less negative than in the C- and D-zones (Fig. 1D, Table S2), indicating a more disordered motor orientation in the P-zone, where myosin interacts with the thin filament tips.

The space-averaged <*P*_2_> values for the TnC-C and TnC-E probes also decreased in rigor myofibrils compared with their relaxed values, as observed in muscle fibres (Table S1). However, the orientation distributions resolved by FPM revealed that <*P*_2_> for each probe was much lower in the A-band than in the I-band (Fig. 1C, cyan and magenta open circles; Fig. 1D). These results show that myosin motors bound to actin in rigor trigger large structural changes in troponin within the filament overlap region in the A-band, which are partially propagated along the thin filament into the I-band.

In the presence of calcium ions (pCa 4.7) the RLC orientation distribution in rigor myofibrils remained unchanged (Fig. 1C, black circles). By contrast, the orientation distribution of the TnC-C probe shifted towards more negative values (Fig. 1C, cyan filled circles), consistent with a similar reduction of <*P*_2_> in both I- and A-bands (Fig. 1D). Instead, <*P*_2_> for the TnC-E probe decreased predominantly in the A-band, without significant changes in the I-band (Fig. 1C, magenta filled circles; Fig. 1D). These results demonstrate that, in addition to the myosin-induced structural changes in troponin, calcium induces further structural changes in the regulatory head of troponin (detected by the TnC-C probe) that are additive along the entire length of the thin filament. By contrast, calcium-induced changes in the IT-arm of troponin (detected by the TnC-E probe) are additive mainly within the region of filament overlap. Taken together, these findings indicate that, under rigor conditions at saturating calcium and in the absence of filament force, thin filament activation is markedly enhanced within the filament overlap region (A-band) compared to the non-overlap region (I-band), highlighting a spatially non-uniform regulatory state of the thin filament induced by the attachment of myosin motors to actin.

### The folded-OFF state of myosin motors is enriched in the C-zone of the thick filament in relaxed myofibrils at physiological temperature and lattice spacing

Since the conformation of myosin motors in relaxed muscle fibres is sensitive to temperature (28), we investigated the effect of increasing the temperature in the range 12-30°C on the spatial distribution of RLC probe orientations in relaxed myofibrils. The temperature increase induced an increase in the spatially-averaged <*P*_2_> for the RLC-E probe in the sarcomere of a myofibril similar to that measured by FISS in relaxed muscle fibres (28) (Fig. S3), consistent with an increase in the fraction of folded motors with parallel orientation with respect to the filament axis. Instead, both TnC probes were much less sensitive to the temperature increase (Fig. S4), as observed in muscle fibres (12).

The spatial distribution of RLC probe orientations resolved by FPM showed that, at each temperature, the <*P*_2_> values in the P- and C-zones of the thick filament were higher than those in the D-zone (Fig. 2A). Indeed, our model simulations (Fig. 2A, solid lines) show that the temperature-induced increase in <*P*_2_> was greater in the P- and C-zones than in the D-zone, and, at temperatures above 27 °C, <*P*_2_> reached its maximum in the C-zone (Fig. 2B). These findings indicate that at near-physiological temperature the fraction of folded OFF myosin motors is higher in the C-zone.

**Figure 2.**
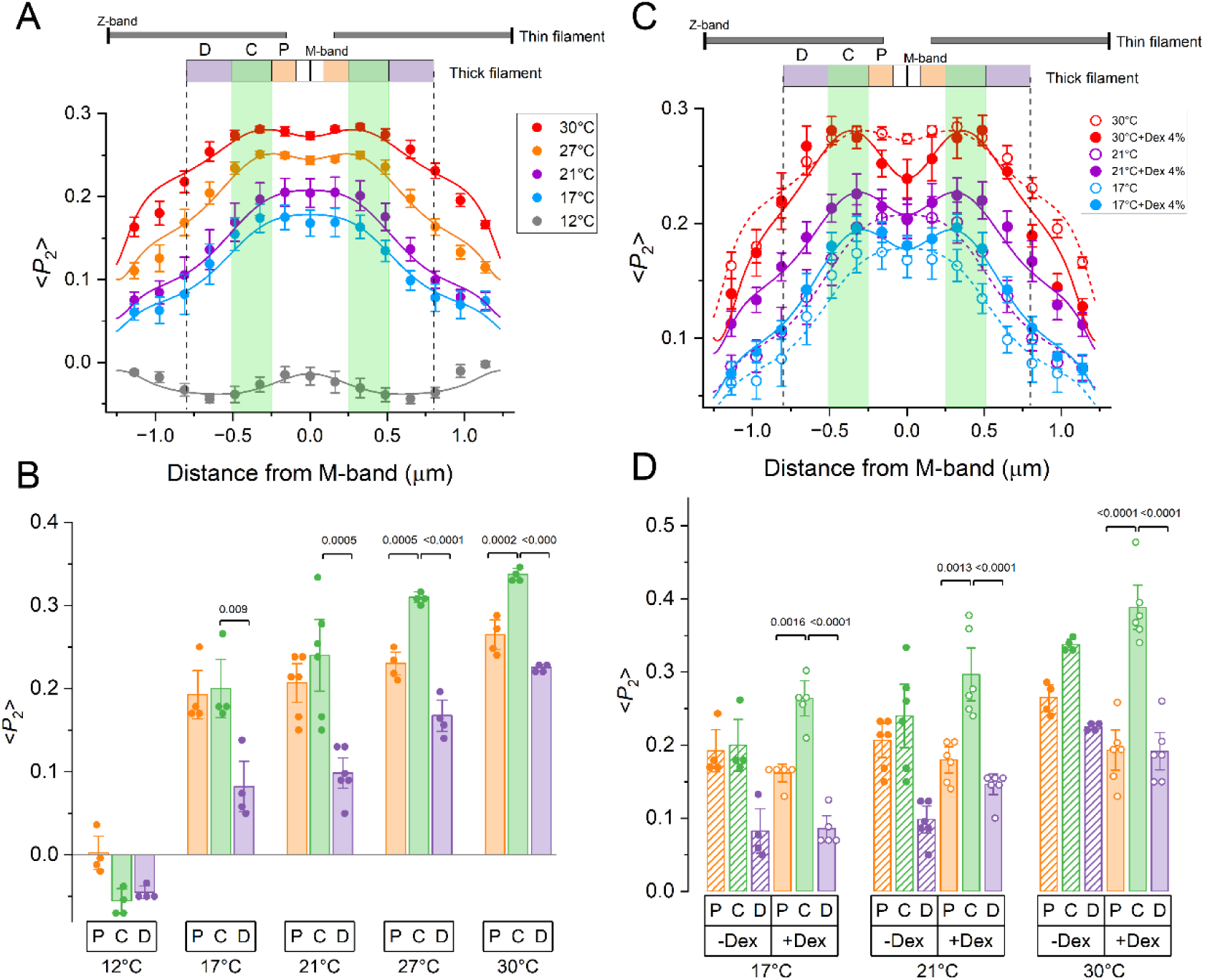
Effect of temperature and osmotic compression on the spatial distributions of RLC orientations in relaxed skeletal myofibrils. A) Spatial distribution of <*P*_2_> (mean±SE) for the RLC-E probe along the thick filament in the relaxed sarcomere (pCa 9.0) at 12°C (n=4 myofibrils), 17°C (n=4), 21°C (n=6), 27°C (n=4) and 30°C (n=4), in the absence of Dextran; sarcomere length= 2.55±0.04 µm. Solid lines, simulated distribution of <*P*_2_> for the RLC-E probe using the model described in the Supplementary Text; vertical dashed lines, thick filament tips; green rectangles, C-zone. B) Average <*P*_2_> values (pooled data and mean±SE) in P, C and D zones of the thick filament used to simulate the spatial distribution of <*P*_2_> along the thick filament shown in A. C) Spatial distribution of <*P*_2_> (mean±SE) for the RLC-E probe in the absence (empty circles, same data as in A) and in the presence (filled circles) of 4% (w/v) Dextran T-500, at 17°C (n=5), 21°C (n=6) and 30°C (n=6); sarcomere length= 2.52±0.04 µm. Solid and dashed lines, simulated distribution of <*P*_2_> in the presence and absence of Dextran, respectively. D) Average <*P*_2_> values in P, C and D zones of the thick filament used to simulate the spatial distribution of <*P*_2_> along the thick filament shown in C, in the presence and absence of Dextran.

Next, we tested the effect on RLC orientation of 4% (w/v) Dextran T-500, an osmotic agent known to restore the physiological myofilament lattice spacing and stabilise relaxed structure of the thick filament in demembranated muscle fibres (34). In the presence of Dextran, the cross-sectional area of the myofibril decreased by ∼30%, from 1.49 ± 0.22 µm² (n=40) to 1.07 ± 0.17 µm² (n=34) (mean ± SD), consistent with previously reported changes in both cross-sectional area (35) and lattice cell area measured by X-ray diffraction (34) in demembranated muscle fibres.

TnC probe orientations on the thin filament were mostly unaffected by Dextran (Fig. S4). Instead, at each temperature in the presence of Dextran the spatial distribution of <*P*_2_> for the RLC probe displayed two peaks at ∼±400 nm from the M-band (Fig. 2C, filled circles), coinciding with the centre of the C-zone (36) (Fig. 2C, green bands), generated by higher values of <*P*_2_> for the RLC-E probe in the C-zone than those in the P- and D-zones (Fig. 2D).

Together, these findings indicate that restoring the physiological filament distance in the lattice of the relaxed myofibril specifically enhances the population of folded motors in the C-zone of the thick filament. This strongly supports the hypothesis that MyBP-C crosslinks between thin and thick filaments contribute to stabilisation of the OFF state of myosin within this filament domain.

### Stretch of relaxed myofibrils partially activates the folded-OFF myosin motors in C- and D-zones of the thick filament

To test whether the folded OFF myosin motors are activated by filament stress independently of calcium, we studied the changes in RLC orientation along the thick filament induced by stretching the relaxed myofibril in near physiological conditions (T=30°C, 4% w/v Dextran T-500) (Fig. 3). The relationship between passive tension and sarcomere length in single myofibrils was similar to that previously reported (37) and was not affected by the introduction of the TnC or RLC probes (Fig. 3B).

**Figure 3.**
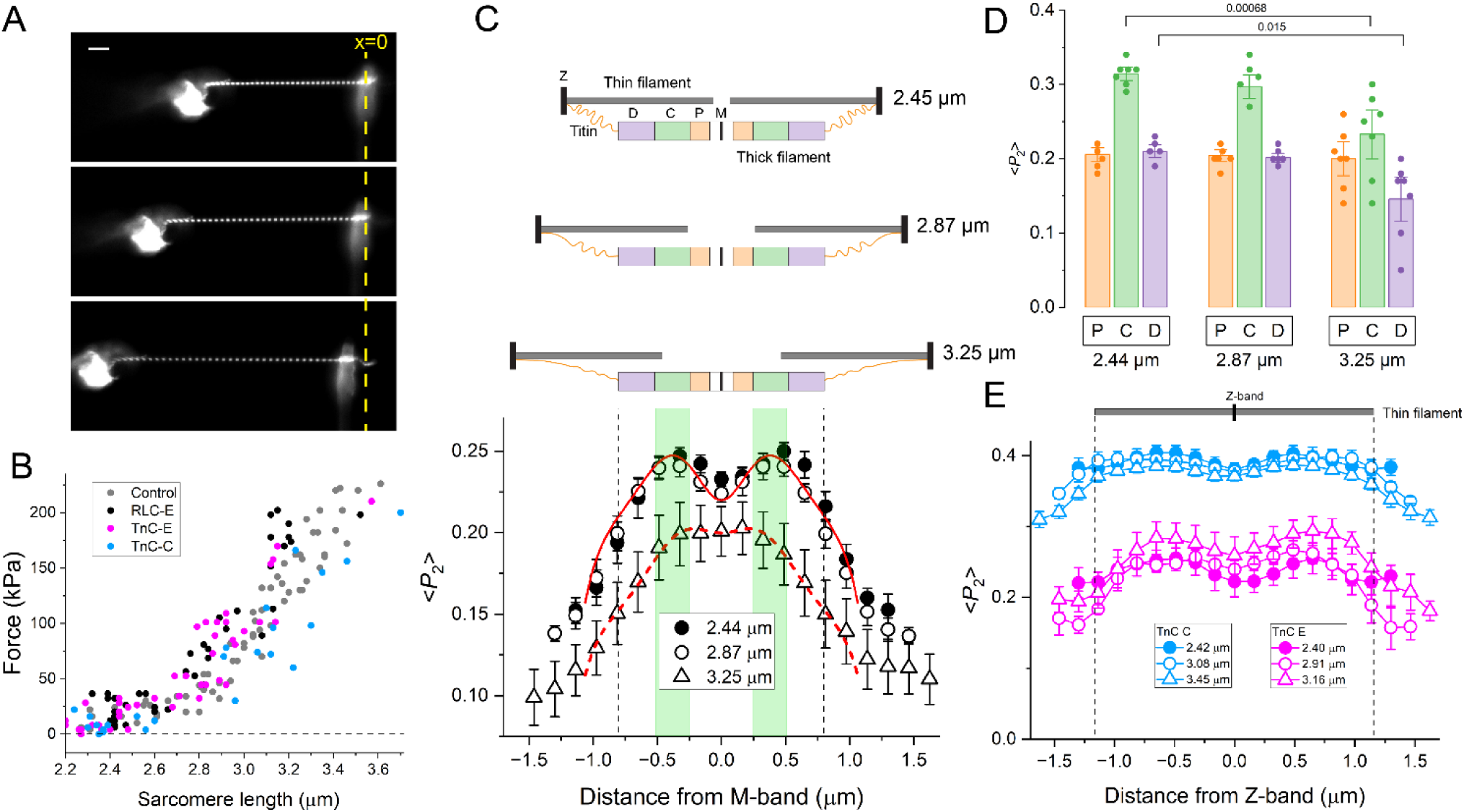
Sarcomere length-dependent activation of the myosin motors in C and D zones of the thick filament in relaxed skeletal myofibrils. A) Fluorescent images of a single skeletal myofibril exchanged with the RLC-E probe mounted between the puller (left) and cantilever (right) in relaxing solution (pCa 9.0) at the initial resting length (top) and after stretches of 16% (middle) and 32% (bottom) of the initial length. Temperature=30°C, 4% (w/v) Dextran T-500. The yellow dashed-line marks the resting position of the cantilever (zero force). Scale bar, 10 µm. B) Passive force-sarcomere length relationship in single myofibrils without probes (Control, grey; n=8 myofibrils) and myofibrils exchanged with RLC-E (black; n=10), TnC-E (magenta; n=14) and TnC-C (cyan; n=10) probes. C) Top panels, schematics of the sarcomere at the three myofibril lengths in (A). Lower panel, average distributions of <*P*_2_> (mean±SE; n=7) for the RLC-E probe along the thick filament in the relaxed sarcomere (pCa 9.0) at the three sarcomere lengths. Solid and dashed red lines, simulated <*P*_2_> distributions for the RLC-E probe along the thick filament at short and long sarcomere lengths; dashed vertical lines, thick filament tips; green bands, thick filament C-zone. D) Average <*P*_2_> values (pooled data and mean±SE) in P-, C- and D-zones of the thick filament used to simulate the spatial distribution of <*P*_2_> along the thick filament in each myofibril at the three sarcomere lengths. E) Sarcomere-length dependence of <*P*_2_> distributions (mean±SE) along the thin filament for the TnC-C (n=7) and TnC-E (n=5) probes in relaxed myofibrils.

Increasing the sarcomere length from 2.2 to 2.8 µm slightly increased the passive tension of the myofibril to 50 kPa (Fig. 3B), corresponding to ∼10% of the isometric force (*T*_0_) during activation at maximal calcium concentration (Fig. 4A). At this sarcomere length the overlap of the thin filament with the P-zone of the thick filament was abolished (Fig. 3C), but the spatial distributions of RLC (Fig. 3C, open circles) and TnC probe orientations (Fig. 3E) were unaffected.

**Figure 4.**
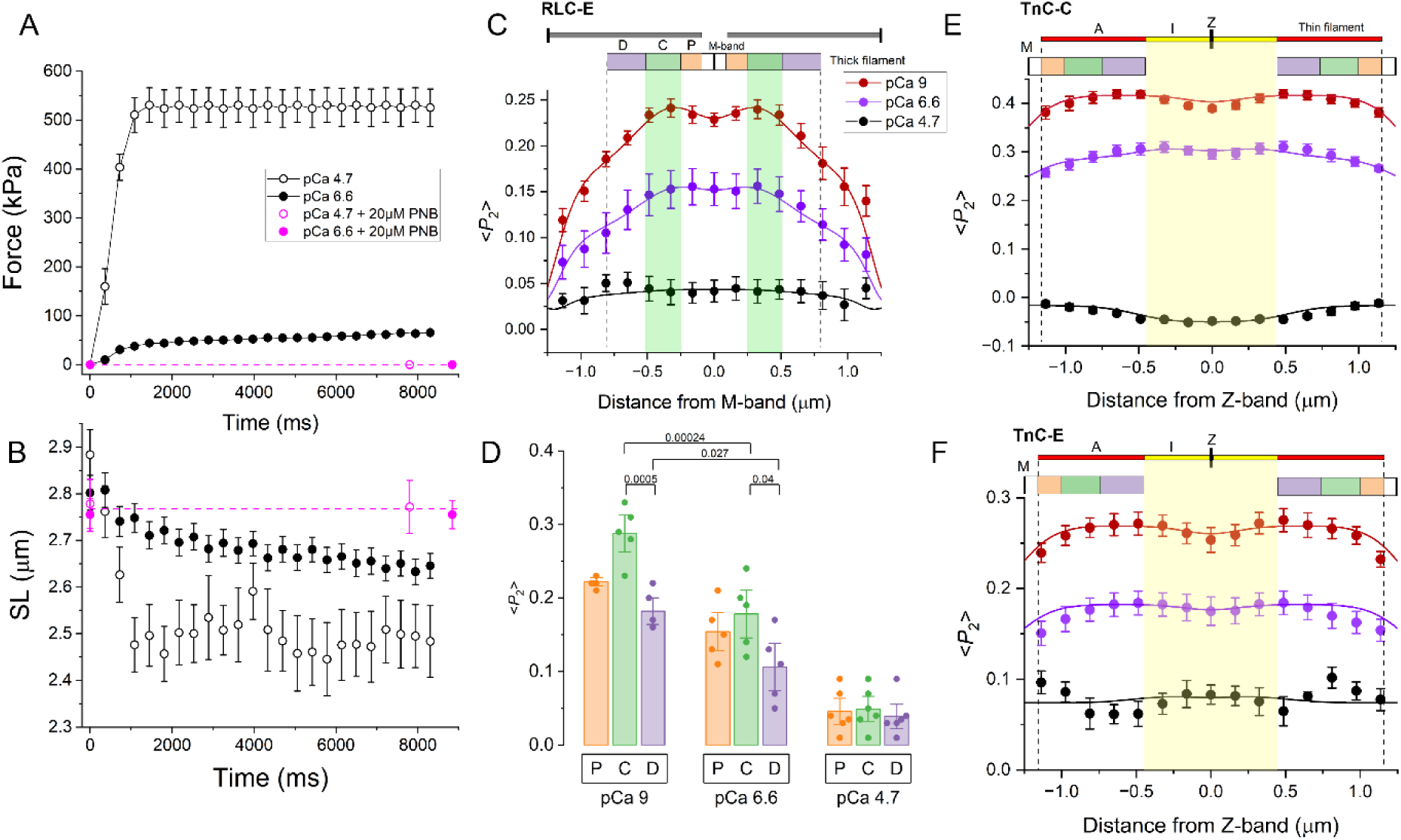
Spatial distributions of RLC and TnC orientations in the sarcomere of Ca^2+^-activated myofibrils. A) Time course of force and B) sarcomere length (SL) in single skeletal myofibrils activated at time zero at pCa=6.6 (filled symbols) and pCa=4.7 (empty symbols), in the absence (black, mean±SE, n=15) and in the presence (magenta, mean±SE, n=17) of 20 µM Para-Nitro Blebbistatin (PNB). Temperature=30°C. Magenta dashed lines mark the zero force and the average SL before activation in the presence of PNB. C) Spatial distribution of <*P*_2_> for the RLC-E probe along the thick filament (circles) and model simulations (lines) before activation (pCa 9) and at the steady state of activation at pCa 6.6 and 4.7 (n=6 myofibrils for pCa 4.7, n=5 for RLC-E for pCa 9 and 6.6). D) <*P*_2_> values for the RLC-E probe (pooled data and mean±SE) in P-, C- and D-zones of the thick filament used to simulate the spatial distribution of <*P*_2_> in C. E,F) Spatial distributions of <*P*_2_> for TnC-C and TnC-E probes along thin filaments (circles) and model simulations (lines) at pCa 9, 6.6 and 4.7 (n=6 myofibrils). Model parameters are shown in Table S3. Yellow square, I-band at 2.5 µm sarcomere length.

Stretching the myofibril to sarcomere lengths higher than 2.8 µm induced a sharp increase in passive tension (Fig. 3B) due to the stretch of the titin links between the thick filament tips and the Z-band (Fig. 3C). At a sarcomere length of 3.3 µm the overlap between the thin filament and the C-zone was almost abolished and the titin-based passive tension increased to 150-200kPa (∼30% of *T*_0_) (Fig. 3B,C). The orientation of the TnC probes on the thin filament remained constant in the whole sarcomere length range analysed (Fig. 3E), indicating that the regulatory state of the thin filament was not affected by stretch. In contrast, <*P*_2_> for the RLC-E probe decreased and the two C-zone peaks characteristic of the <*P*_2_> distribution at shorter sarcomere lengths disappeared (Fig. 3C, open triangles).

Modelling of the RLC orientation along the thick filament showed that at 3.3 µm sarcomere length the average <*P*_2_> for the RLC-E probe decreased in the C- and D-zones of the thick filament whereas that in the P-zone did not change significantly (Fig. 3D). These results show that stretching the myofibrils in the absence of calcium at sarcomere lengths longer than 2.8 µm induce more perpendicular orientations of myosin motors in the C- and D-zones of the thick filament, consistent with a reduction in the fraction of folded-OFF motors in these filament domains.

### Spatial control of thin- and thick-filament activation in calcium-activated myofibrils

To study the activation of troponin and myosin in regulatory domains of thin and thick filament, respectively, we performed FPM measurements of TnC and RLC probe orientations in myofibrils activated by calcium at near-physiological temperature (30°C). At saturating calcium concentration (pCa=4.7) the force of the myofibril increased up to ∼530 kPa (Fig. 4A), as previously reported (37). We monitored changes in the orientation of TnC and RLC probes in a population of sarcomeres in the central region of the myofibril which shortened from 2.9 to 2.4 µm (Fig. 4B, open symbols). In relaxed myofibrils, <*P*_2_> for the RLC-E probe was higher in the C-zone, but it markedly decreased upon activation at pCa 4.7 across all filament domains, becoming uniform along the thick filament (Fig. 4C,D), signalling the release of myosin motors from the folded state throughout the filament. <*P*_2_> for the TnC-C and E-probes also decreased to values that were uniform along the thin filament (Fig. 4E,F; Table S3), indicating a homogenous activation of the thin filament in both I- and A-bands.

Activation of myofibrils at submaximal calcium concentration (pCa 6.6) increased active force in the myofibril to ∼70 kPa, corresponding to ∼15% of the force at maximal calcium, and was accompanied by sarcomere shortening from 2.8 to 2.6 µm (Fig. 4A,B; filled circles). The spatial distribution of <*P*_2_> for the RLC-E probe was shifted to intermediate values between the relaxed state (pCa 9) and the fully activated state (pCa 4.7) (Fig. 4C, purple circles; Fig. S6), and was simulated with higher <*P*_2_> values in the P- and C-zones than in the D-zone (Fig. 4D; Table S3). These results indicate that, at submaximal [Ca²⁺] when thick filament activation is half-maximal, myosin motors in the P- and C-zones remain preferentially stabilised in the OFF state, whereas motors in the D-zone show a greater degree of activation. In contrast, the thin filament exhibited a uniform activation level in I and A-bands, as shown by the homogenous values of <*P*_2_> along the thin filament (Fig. 4E,F; purple circles; Table S3).

To determine the relative contribution of calcium and myosin to the activation of the thin filament we activated the myofibrils in the presence of the myosin inhibitor Para-Nitro-Blebbistatin (PNB) (38). At 20 µM, PNB completely inhibited force generation and sarcomere shortening at both activating calcium concentrations (Fig. 4A,B; magenta circle), and abolished the associated changes in RLC orientation (Fig. 5A), thereby stabilizing the myosin motors in a spatial distribution of OFF conformations along the thick filament, similar to that observed in relaxed myofibrils in the absence of PNB (Table S3). The calcium-induced changes in <*P*_2_> for the TnC-C probe were largely preserved at both calcium concentrations in the presence of PNB, although at pCa 4.7 <*P*_2_> in the I-band was higher than that in the absence of PNB (Fig. 5B; Table S3). In contrast, PNB largely abolished the orientation changes of the TnC-E probe in both the I- and A-bands, such that at maximal calcium activation (pCa 4.7) the amplitude of the <*P*_2_> change was reduced to ∼25% of that observed during contraction in the absence of PNB in both filament domains (Fig. 5C; Table S3). These results indicate that the orientation of the regulatory head of troponin, reported by the TnC-C probe, depends primarily on calcium binding to TnC in the A-band, with only a minor contribution from myosin in the I-band. Conversely, the orientation of the IT-arm of troponin along the thin filament, reported by the TnC-E probe, is modulated by both calcium and myosin, with myosin accounting for ∼75% of the structural change observed during maximal calcium activation.

**Figure 5.**
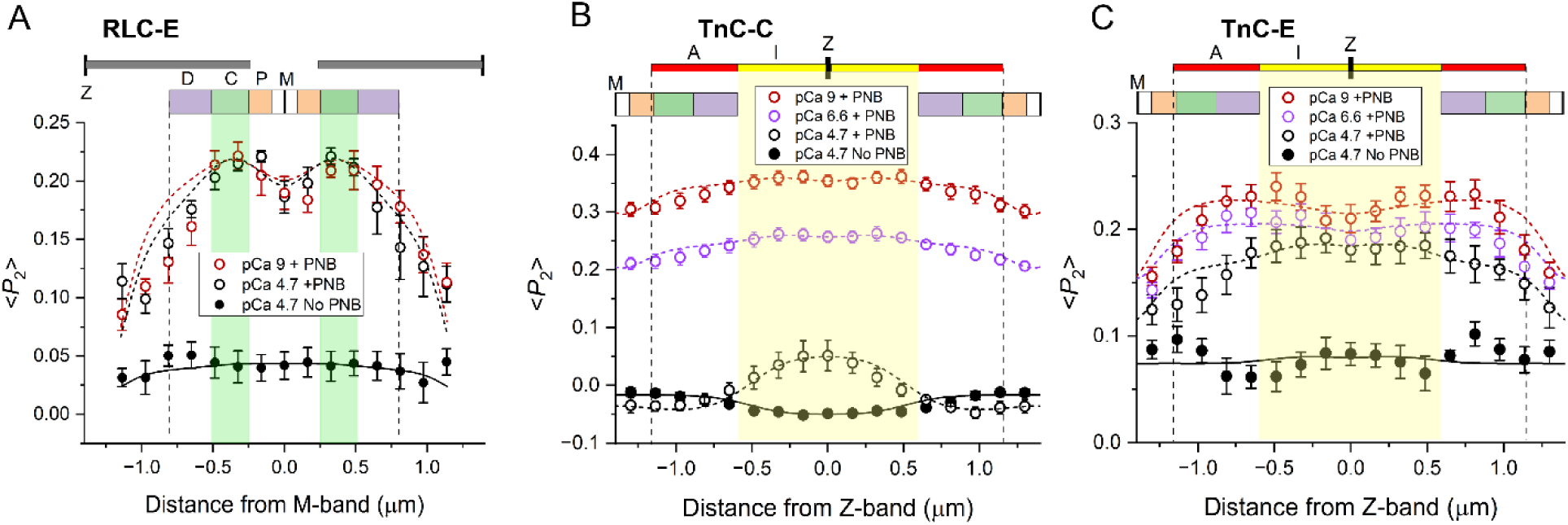
Effect of PNB on the spatial distribution of RLC and TnC probes in the sarcomere of calcium-activated myofibrils. Spatial distributions of <*P*_2_> for RLC-E (A), TnC-C (B) and TnC-E (C) probes at pCa 9/6.6/4.7 in the presence of the myosin inhibitor PNB (20 µM) (n=6 myofibrils for each probe) and at pCa 4.7 in the absence of PNB (filled black circles, as shown in Fig. 4). Yellow squares, I-band width at 2.8µm sarcomere length. Short-dashed lines, model simulations of probe orientation distribution in the presence of PNB. Solid black lines, model simulations of data in the absence of PNB at pCa 4.7. Model parameters are shown in Table S3.

## Discussion

Here, we introduce fluorescence polarization microscopy (FPM) as a novel approach to resolve the regulatory states of both thin and thick filaments within distinct sarcomere domains in isolated myofibrils under near-physiological conditions. Space-averaged orientations of the myosin regulatory light chain (RLC) and troponin-C (TnC) measured by FPM closely matched bulk values obtained by FISS in muscle fibres, confirming that the physiological regulatory transitions in the myofilaments are preserved in isolated myofibrils. Additionally, our FPM method allowed us to resolve myosin and troponin regulation within single sarcomeres, uncovering details inaccessible to fibre-averaged analyses.

Using FPM with a probe on the RLC, we first mapped the distribution of OFF/ON myosin motors across the P-, C-, and D-zones of the thick filament in relaxed myofibrils. Increasing temperature induced a non-uniform increase in the fraction of folded-OFF motors in thick filament domains, with the highest fraction in the C-zone at physiological temperature (Fig. 2), providing direct in-situ evidence that MyBP-C stabilizes the folded-OFF myosin state in this filament domain. Restoration of native lattice spacing by osmotic compression further stabilized this folded conformation in the C-zone, strongly suggesting that the stabilization of the myosin OFF states requires the formation of N-terminal links of MyBP-C with actin (Fig. 6A), in addition to the C-terminal interaction with myosin heads on the thick filament (18, 19). This conclusion is consistent with the evidence that cleavage of N-terminal domains of MyBP-C promotes the release of myosin motors from the OFF state (39).

**Figure 6.**
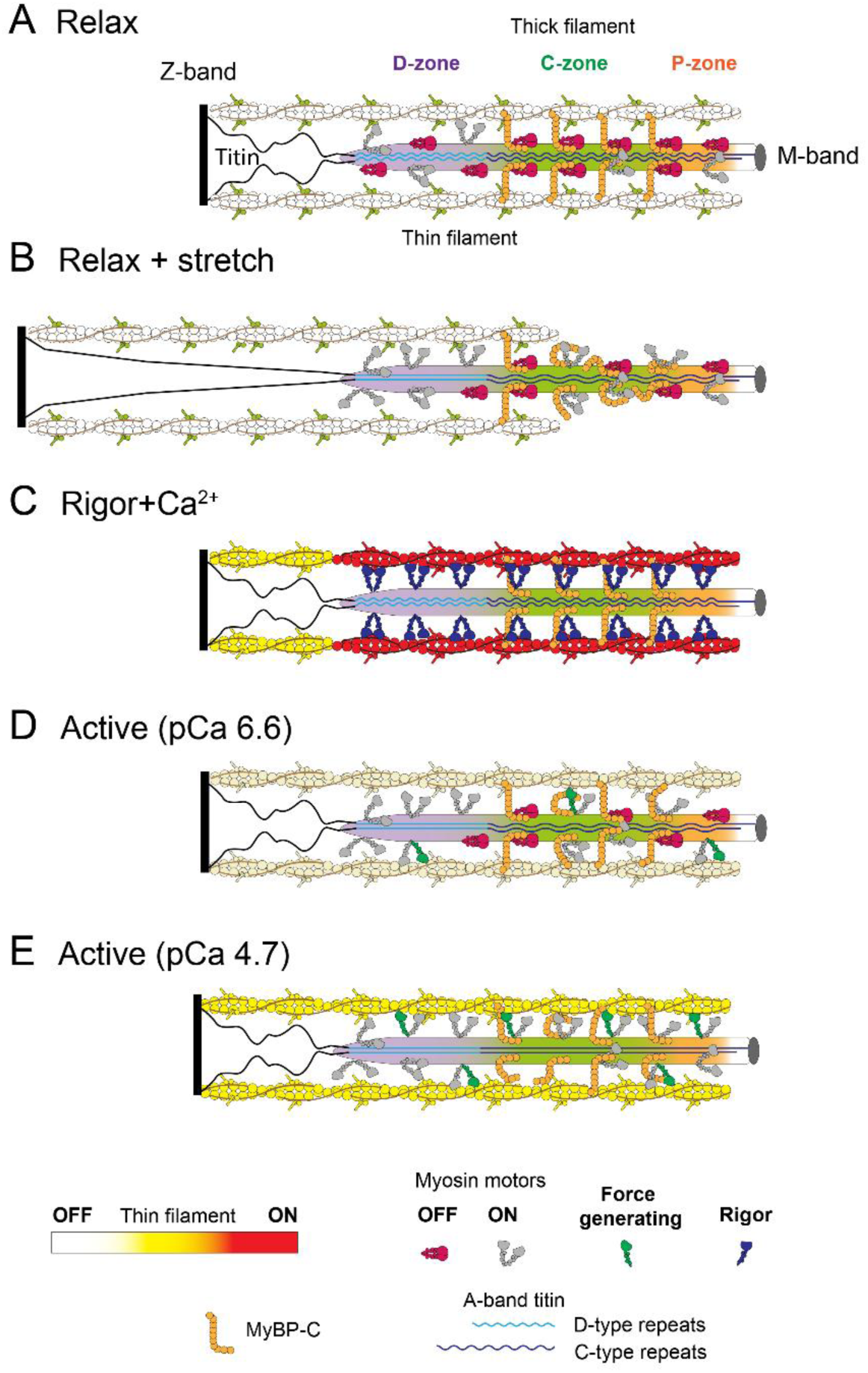
Zonal activation of thin and thick filaments in the sarcomere of skeletal muscle. Schematic of myosin motor conformations in P, C, D zones of the thick filament and thin filament activation in I-and A-bands in the half-sarcomere in relaxed, rigor and calcium-activated myofibrils. A) In the absence of calcium (pCa 9) and under zero filament force the thin filament is in the blocked-OFF state, and the folded-OFF state of myosin is enhanced in the C-zone of the thick filament. Constitutively-ON motors are shown in grey. B) Stretching of the titin links at long sarcomere lengths (≥3.3µm, pCa 9) causes an increase in the passive force applied on the thick filament associated with conformational changes in D-type and C-type titin super-repeats, in D- and C-zones respectively, and loss of MyBP-C links in the C-zone. Folded myosin motors are released from the OFF state in D- and C-zones. C) Under rigor conditions and zero filament force in the presence of calcium, myosin motors are strongly bound to actin, inducing the transition of the thin filament to the Open state (red) in the A-band and to a partially activated state (yellow) in the I-band. D) At intermediate calcium concentrations (pCa 6.6) and submaximal force the thin filament is uniformly activated along its length, whereas in the thick filament the folded-OFF motors are mainly located in the P- and C-zones. E) At maximal calcium (pCa 4.7) and higher filament forces, both thin and thick filaments are uniformly switched ON.

We next asked whether the distribution of OFF myosin motors resolved by FPM corresponds to that of the energy-saving SRX state of myosin identified in spatially resolved measurements of ATP turnover in relaxed myofibrils. These studies reported a concentration of SRX motors in the C-zone (21, 22), consistent with the enrichment of OFF motors in the same region observed here by FPM. However, despite the strong temperature-dependence of the myosin OFF state (Fig. 2), the SRX state of myosin in myofibrils is largely temperature-insensitive (40), clearly highlighting the mismatch between structural and biochemical states of myosin. Thus, our results support the emerging view that the folded-OFF structural state of myosin does not directly correspond to the SRX biochemical state (23, 25), implying that distinct myosin subpopulations collectively account for the slow ATP turnover observed in relaxed skeletal muscle.

To test whether folded motors can be activated by mechanical stress independently of calcium, we examined stretch-induced structural changes in relaxed myofibrils. Increasing sarcomere length beyond 2.8 μm induced an increase in titin-based passive tension and a more perpendicular orientation of myosin motors on the thick filament, consistent with the stress-dependent activation of the thick filament in relaxed muscle fibres stretched at longer sarcomere length (35, 41). However, our FPM analysis showed that the fraction of perpendicular-ON motors increased in the C- and D-zones, but not in the P-zone. Activation of OFF motors in the D-zone may arise from structural rearrangements of D-type titin super-repeats along the thick filament backbone under tension, whereas in the C-zone, partial activation likely results from the disruption of MyBP-C links due to reduced filament overlap at longer sarcomere lengths (Fig. 6B).

We next analyzed the spatial distribution of active myosin motors across thick-filament zones during contraction. At maximal calcium levels (pCa 4.7), myosin motors were fully activated throughout the filament (Fig. 6E). In contrast, at submaximal calcium (pCa 6.6), FPM revealed a gradient of myosin activation, with a greater proportion of folded-OFF motors persisting in the P- and C-zones (Fig. 6D), despite these regions experiencing higher filament stress than the D-zone. Therefore, these results point to a larger stability of the myosin OFF state in P- and C-zones, and support a zonal model of thick filament activation, in which D-zone motors are activated at lower stress, whereas C- and P-zone motors require higher filament tension for full activation.

By using FPM with TnC probes on the regulatory head (TnC-C) and IT-arm (TnC-E) of troponin we resolved the zonal regulation of thin filaments within A- and I-bands of the sarcomere. In relaxed myofibrils both TnC probes showed a uniform orientation, consistent with the fully OFF or Blocked state of the thin filament (Fig. 6A). We then isolated the effect of myosin attachment to actin on thin filament activation in rigor myofibrils in the absence of force.

Under these conditions, myosin motors induced a greater activation of the thin filament in the A-band than in the I-band, consistent with cryo-ET reconstructions showing greater tropomyosin displacement in the A-band (42). Addition of calcium produced additive structural changes in the regulatory head of troponin along the entire filament, but in the IT-arm these changes were confined to the A-band (Fig. 1). We therefore conclude that in rigor myofibrils, in the absence of force, full activation of the thin filament by myosin and calcium occurs only within the A-band, corresponding to the Open state described in in vitro studies of isolated filaments (43), and that the myosin-dependent structural changes in the thin filament are partially transmitted to the I-band, where acto-myosin interactions are absent (Fig. 6C).

To isolate the effect of calcium on thin filament regulation, we analysed probe orientations at saturating calcium concentrations in the presence of the myosin inhibitor PNB, which prevents motor attachment to actin and force generation (38). Under these conditions, calcium-induced orientation changes in the regulatory head of troponin within the I-band were comparable to those observed in rigor at pCa 4.7, while those in the A-band were slightly reduced (Fig. S7). These findings indicate that at saturating calcium and at physiological concentrations of ATP the troponin regulatory head becomes largely independent of myosin binding, as previously observed in intact fibres (12). In contrast, the amplitude of the calcium-induced structural changes in the IT-arm was only ∼25% and ∼10% of that observed in rigor + Ca conditions in I- and A-bands, respectively (Fig. S7). Since tilting of the IT-arm is coupled with the azimuthal shift of tropomyosin on actin (4), we conclude that calcium alone elicits only a partial activation of the thin filament, corresponding to the shift from the Blocked to the Closed state (43).

We next examined how these regulatory mechanisms operate during physiological isometric contraction, when the thin filament is mechanically loaded. Unexpectedly, at maximal calcium activation, the orientation of both troponin regulatory head and IT-arm was uniform along the thin filament (Fig. 4), in contrast with the dual regulatory states observed in the I-and A-bands of rigor myofibrils when filaments are unloaded. While the orientation change of the troponin regulatory head was primarily driven by calcium, the amplitude of the IT-arm orientation change in both the I- and A-bands was approximately fourfold greater than that induced by calcium alone (Table S3), indicating that thin filament activation, as reported by the IT-arm, remains predominantly dependent on myosin attachment even under physiological conditions. This observation agrees with the conclusion of a previous study that the myosin-dependent component accounts for ∼70% of the orientation change of the TnC-E probe in maximally activated muscle fibres (12), derived from bulk orientation measurements assuming no propagation of activation from the A- to the I-band. Using FPM, we directly resolved these myosin-dependent structural changes in both thin filament domains during contraction. The amplitude of the IT-arm orientation change in the A-band at maximal activation (pCa 4.7) was ∼30% of that observed in rigor + Ca (Fig. S7), consistent with the lower fraction (∼30%) of actin-attached motors during isometric contraction at maximal calcium (44). This finding indicates that the level of activation of the thin filament in the A-band primarily depends on the number of force-generating motors attached to actin.

However, the fraction of actin-attached motors during a physiological isometric contraction is insufficient to induce a complete transition of the thin filament to the Open state in the A-band (Fig. 6E). Notably, the orientation of the IT-arm in the I-band during contraction was similar to that in rigor (Fig. S7), despite fewer actin-attached motors, and matched the values observed in the A-band at both low and high calcium concentrations. We therefore propose that during contraction, when the thin filament is mechanically loaded, myosin-dependent regulatory structural changes in the thin filament initiated in the A-band are transmitted to the I-band -where actin–myosin interactions are absent but thin filament tension is maximal-via a force-dependent mechanism that ensures coordinated activation of the thin filament across sarcomeric domains (Fig. 6D,E).

All together, these results establish FPM as a powerful method for visualizing zone-specific regulatory transitions of myofilaments under near-physiological conditions. By demonstrating that the activation of both thin and thick filaments is spatially heterogeneous and mechanically modulated, this work defines a dual-filament regulatory paradigm of muscle contraction at sub-sarcomere level that provides a structural framework for investigating muscle function, disease, and targeted therapeutic modulation.

## Methods

### Preparation of TnC- and RLC-labelled skeletal myofibrils

Muscle bundles (length ∼4 cm, width ∼2 mm) were dissected from the psoas muscle of adult (18 weeks old) New Zealand White rabbits. The rabbits were sacrificed by injection of an overdose of sodium pentobarbitone (200 mg kg^-1^) in the marginal ear vein, followed by confirmation method, in agreement with Schedule 1 of the Animals (Scientific Procedures) Act 1986 (UK). A total of six rabbits were used in this study. The bundles were stored in rigor solution (28) containing 50% (vol/vol) glycerol at −20°C for at least 1 week before the experiments. On the day of the experiment smaller bundles (length ∼3 mm, width ∼1 mm) were dissected and transferred to rigor solution at ∼1°C for 10 min, then washed in RLC/TnC Extracting Solution (20 mM EDTA, 50 mM Potassium Propionate, 10 mM Potassium Phosphate buffer, pH 7.1) at ∼1°C for 10 min. About 30% of the native RLC/TnC was exchanged with labelled proteins by transferring the bundles into Extracting Solution containing ∼20 µM of labelled RLC/TnC and incubated for 1h at room temperature, as previously described (28). The preparation of the TnC variants labelled with bifunctional rhodamine (BR) on the E-helix (E96C/R103C; TnC-E probe) and C-helix (E56C/E63C TnC-C probe), and of the RLC variant labelled with a bifunctional sulphorhodamine (BSR, B10621, Invitrogen) on the E-helix (D95C/V103C; RLC-E probe), as well as the composition of the physiological buffers, were described previously (12, 28, 29). For the rigor experiments, the bundles were washed three times in rigor solution for 10 min to wash out any unbound protein, and homogenised into 1.2 ml of rigor solution using an Ultra-Turrax disperser T10 with S10N-5G dispersing element (IKA, UK) for 20s at speed 5, followed by 20s at speed 4. For all the other experiments the bundles were incubated for 1h at room temperature in relaxing buffer containing wild-type TnC, washed in relaxing buffer and then homogenised in relaxing buffer as described above. The suspension of labelled myofibrils was stored at 4°C up to three days.

### Fluorescence polarization microscopy (FPM) on myofibrils

Single or bundles of 2-3 myofibrils (∼100µm long, 1-3µm wide) were mounted in a temperature-regulated trough in rigor or relaxing solution (28) at pCa=-log[Ca^2+^]=9.0 between two custom-made glass microtools including a puller and a calibrated cantilever force probe (45, 46) (5-20 µm/µN), controlled by two micromanipulators (QUAD, Sutter Instruments, USA) (Fig. S5). The initial sarcomere length was adjusted by stretching the myofibril beyond its slack length. Myofibrils were activated by perfusing them along their length with a stream of activating solution at either maximal (pCa 4.7) or submaximal (pCa 6.6) calcium concentrations. The myofibril setup was mounted on the stage of an inverted microscope (IX73, Olympus) equipped with linear polarisers on a motorised filter wheel (Optospin, Cairn, UK) in the excitation pathway to illuminate the myofibril with linearly polarised light (wavelength 532nm) parallel and perpendicular to the myofibril axis (Fig. S5), similar to a setup previously described (47). The polarization of the excitation light was alternated by switching the position of the linear polarizer in the optical path at a frequency of 2.8Hz. Fluorescence was collected through an objective with magnification limited to 40x and NA=0.7 to reduce depolarization of the fluorescence. Parallel and perpendicular images of the myofibril for each excitation polarization were simultaneously acquired (100-200 ms exposure time) using an image splitter (Optospin II, Cairn, UK) and a CMOS camera (6.5μm x 6.5μm pixel size, 2048 x 2048 array; Prime BSI, Teledyne Photometrics, USA). Using this optical system, we achieved a pixel spacing of 0.162 µm and a spatial resolution (FWHM) of 470 nm (Fig. S 2), which is consistent with the theoretical value for the spatial resolution (SR) of about 440nm with NA=0.7 and wavelength of fluorescence (λ=620nm) (SR=λ/2NA). Following a continuous 10-second exposure of the exchanged myofibril in relaxing conditions, the fluorescence intensity decreased by approximately 5%, without any changes in the order parameter <*P*_2_>, indicating that probe photobleaching was negligible in the experiments presented above.

## Data analysis

The passive and active force generated by the myofibril was determined by measuring the deflection of the calibrated cantilever in the fluorescence polarisation images (Fig. 3A) and normalised to the cross section of the myofibril bundle in the absence of Dextran. Four different fluorescence polarization images of the myofibril (Figure 1 and Fig.S5), ∥I∥, ∥I⊥, ⊥I∥, ⊥I⊥ (where the first and second subscripts indicate the polarization of the excitation and emission, respectively, parallel (∥) or perpendicular (⊥) to the myofibril) were aligned and integrated along the myofibril axis using Image J. The one-dimensional (1-D) polarized intensity profiles were background-subtracted. The second- and fourth-rank order parameters of the orientation distribution of the dipole, <*P*_2_> and <*P*_4_> respectively, and the total fluorescence intensity (*I*) were calculated for each pixel (31), using the values of <*P*_2d_> for each probe, quantifying the rapid probe motion, previously measured in skeletal muscle fibres (12). The average spatial distributions of <*P*_2_>, <*P*_4_> and *I* for each probe in the sarcomere were obtained by averaging the signal distributions in a population of 4-5 adjacent sarcomeres in the myofibril. The average sarcomere length was determined from the total fluorescence intensity distribution of the probe across the same population of sarcomeres.

## Statistical analysis

Differences in <*P*_2_> for RLC-E probe in P, C, D-zones of the thick filament and for TnC probes in A- and I-bands in the protocols shown in Fig. 1-5 were analysed using two-way ANOVA with Tukey’s post hoc analysis. Main effects and post-hoc analyses P-values are provided in the figures and Table Ss 2 and 3. Data presented are mean ± S.E.M. throughout, with (n) indicating the number of myofibrils used. Significance was set at P<0.05 for all analyses.

## Data availability

All data supporting the findings of this study are available within the paper and its Supplementary Information files. Source data for all figures are available at https://doi.org/10.5281/zenodo.17546286 upon publication.

## Acknowledgements

The authors thank Corrado Poggesi and Chiara Tesi (University of Florence, Italy) for providing training in myofibril mechanics in the early stage of the project, and Malcom Irving and Elisabetta Brunello (King’s College London, UK) for their valuable feedback on the manuscript. The authors are grateful to Andras Malnasi-Csizmadia and Anna Rauscher (Motorpharma, Hungary) for generously providing a stock of PNB. This work and the investigators were supported by a Sir Henry Dale Fellowship awarded by the Wellcome Trust and the Royal Society to LF (210464/Z/18/Z). For the purpose of open access, the author has applied a CC BY public copyright licence to any Author Accepted Manuscript version arising from this submission.

## Author contributions

KS and LF contributed to the conception and design of the experiments. LF designed and built the FPM microscope. KS performed the experiments. LF performed model simulations. KS and LF analyzed and contributed to the interpretation of the data. LF wrote the paper. All authors contributed to the critical revision of the manuscript and approved the last version of the article.

