## Supplementary Materials for "Sub-sarcomeric regulation of thin and thick filaments in skeletal muscle myofibrils"

### Supplementary Figures

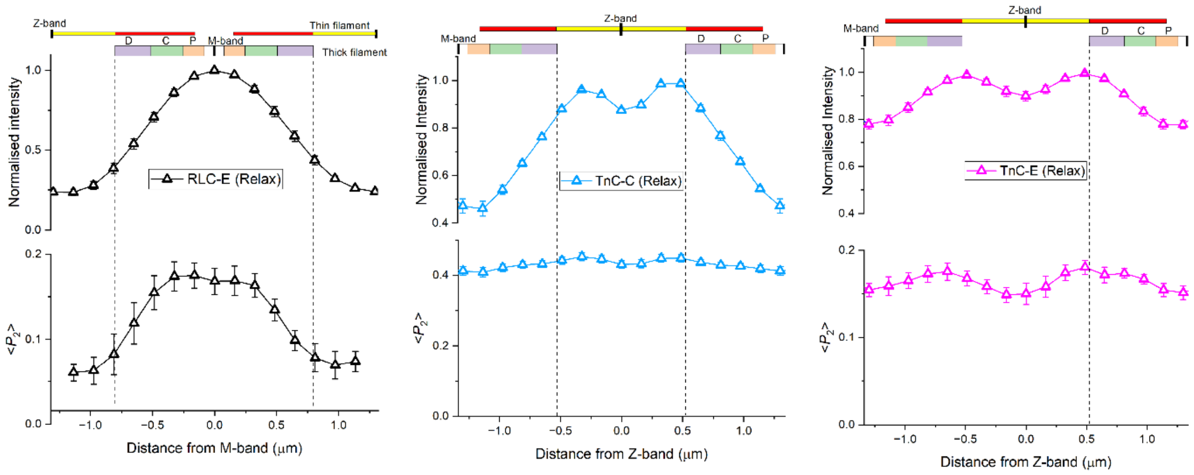


**Fig. S1**. Distribution of Intensity and <$P_2$> for RLC-E probe (black) along the thick filament and for TnC-C (cyan) and TnC-E (magenta) probes along the thin filaments in the sarcomere of relaxed (pCa 9) myofibrils at 17°C.

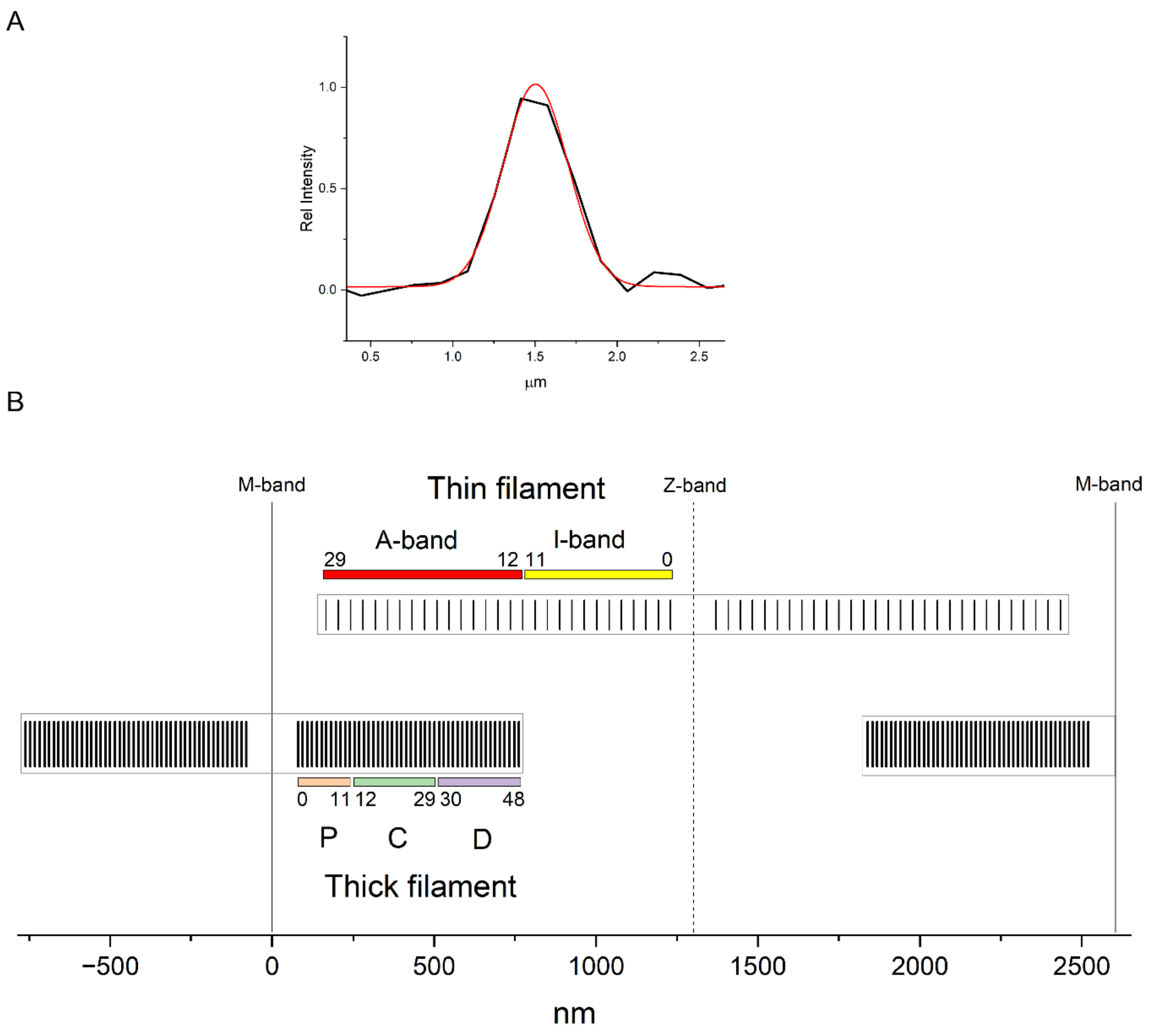


**Fig. S2. A)** Microscope point spread function measured by Gaussian fit (red) of the fluorescence intensity distribution (black) of a point source (fluorescent bead, 170nm diameter). **B)** TnC and RLC probe arrays on thin and thick filaments, respectively, used for the calculation of the spatial distribution of probe orientation in myofibrils (see Supplementary Text). Each probe position on the filaments is marked by small vertical lines. Coloured squares mark P, C, D zones of the thick filaments and A- and I-band domains of the thin filament, with indexes of probe positions at the boundaries of each zone. The sarcomere length measured in the myofibril in each condition was used to set the distance between arrays in adjacent sarcomeres. In the schematic presented above the sarcomere length is 2.6 µm.

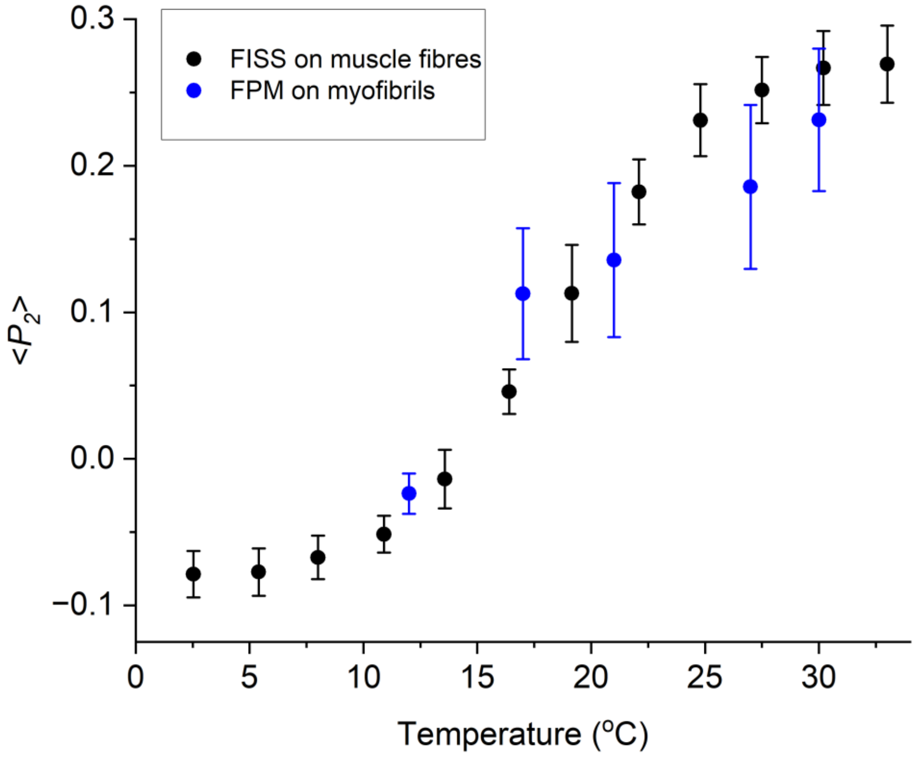


**Fig. S3.** Comparison of the temperature dependence of bulk $<P_2>$ values for the RLC-E probe measured by FISS in demembranated muscle fibres from rabbit psoas (From Fusi et al., 2015)[1] and space-averaged $<P_2>$ measured by FPM in a single sarcomere of skeletal myofibrils from rabbit psoas (Fig.2A) under relaxing conditions (pCa 9) in the absence of Dextran.

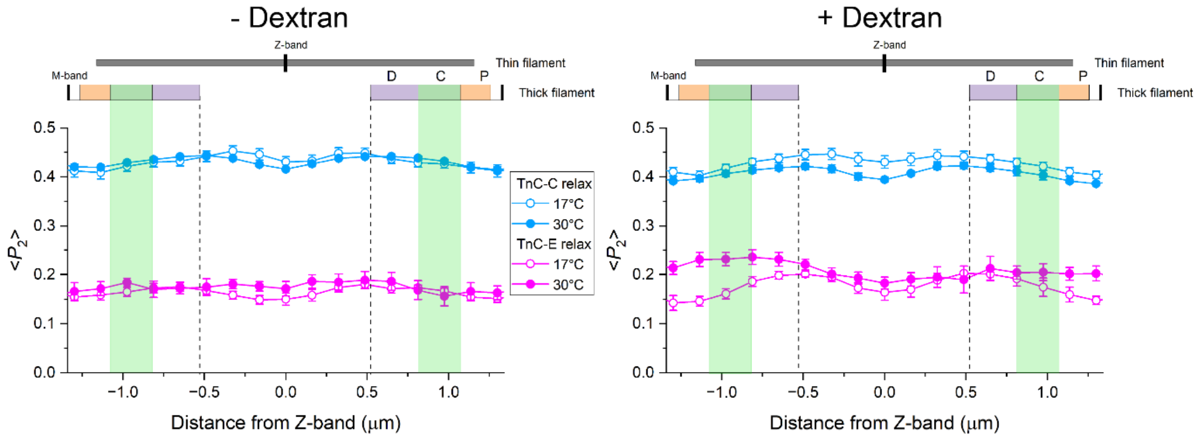


**Fig. S4.** Average distribution of <$P_2$> (mean±SE) for TnC-C (cyan) and TnC-E (magenta) probes along the thin filaments in the relaxed sarcomere (pCa 9.0) of single myofibrils at 17°C (n=5 myofibrils for TnC-C and n=6 for TnC-E) and 30°C (n=5 for TnC-C and n=7 for TnC-E), in the absence and presence of 4% (w/v) Dextran T-500; sarcomere length= 2.52±0.04 µm.

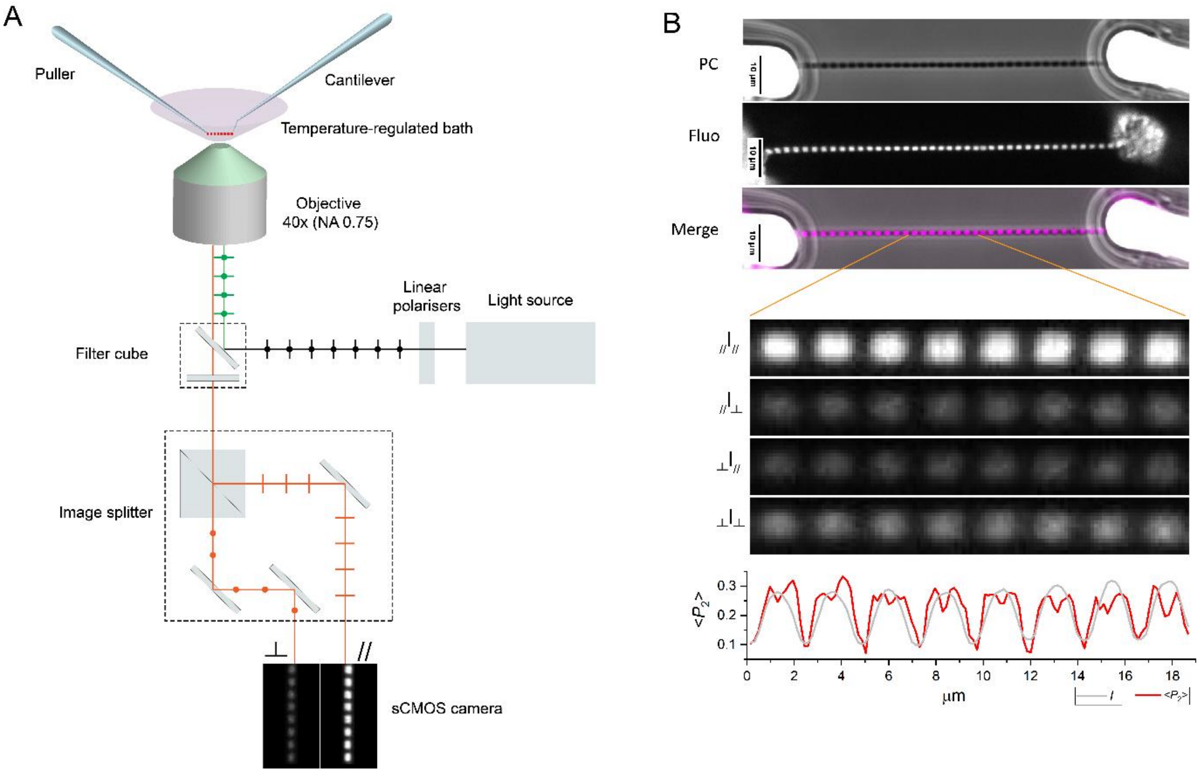


**Fig. S5.** A) Schematic of the Fluorescence Polarization Microscope and myofibril setup. B) Phase contrast (PC) and fluorescence (Fluo) images of a skeletal myofibril from rabbit psoas exchanged with the RLC-E probe in relaxing solution at 30°C. Lower panels, polarised images of sarcomeres in the myofibril (100ms exposure) and the calculated 1D-spatial distribution of <$P_2$> and total fluorescence intensity ($I$) along the myofibril axis (see methods).

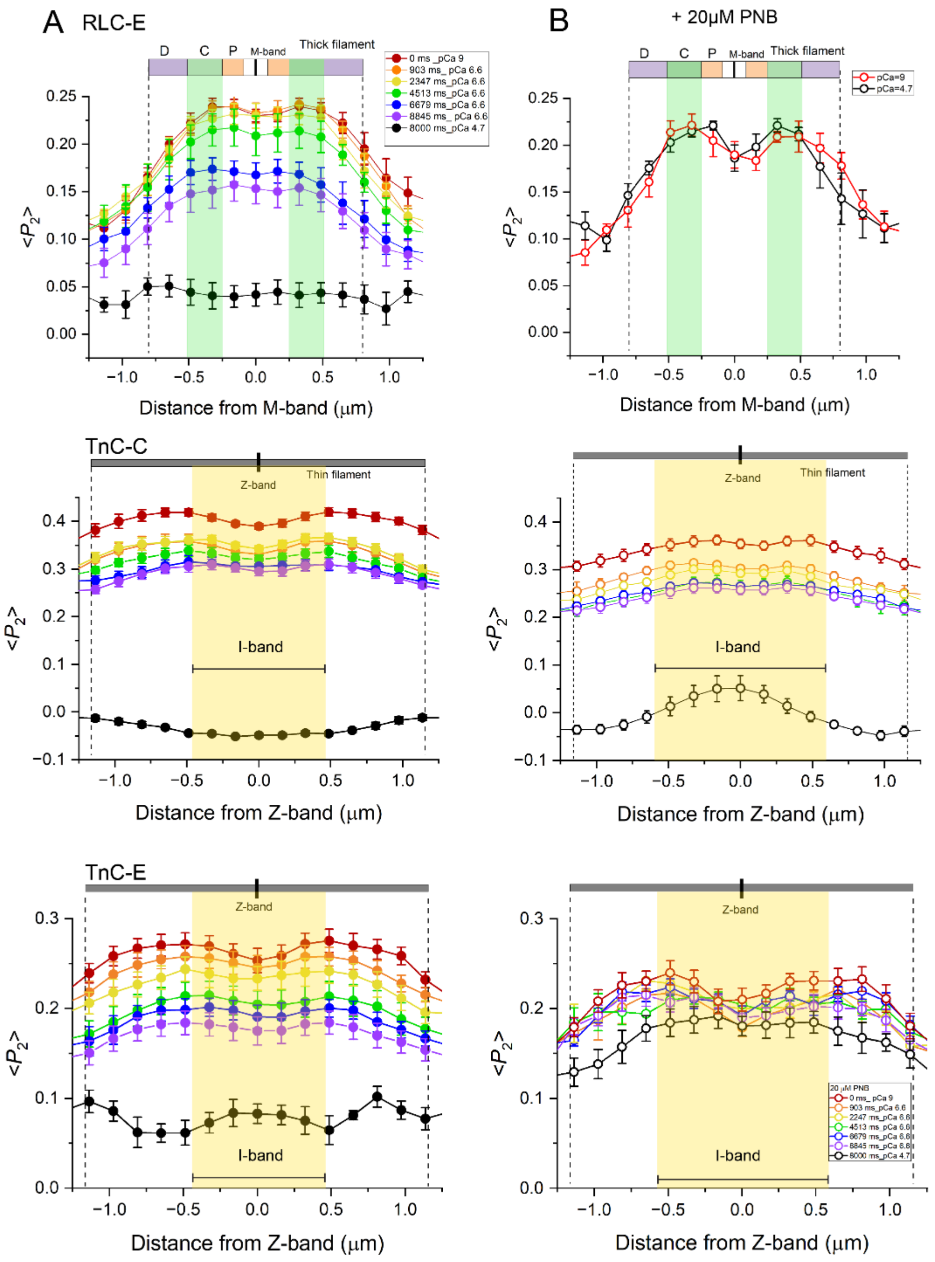


**Fig. S6.** Time courses of the changes in the spatial distributions of <$P_2$> (mean±SE) for RLC and TnC probes along thick and thin filaments, respectively, in myofibril activated at pCa 4.7 and 6.6 at 30°C (Fig. 4) in the absence (left, A) and presence (right, B) of 20 μM Para-Nitro Blebbistatin (PNB). Green squares, thick filament C-zone. Yellow squares, I-band width.

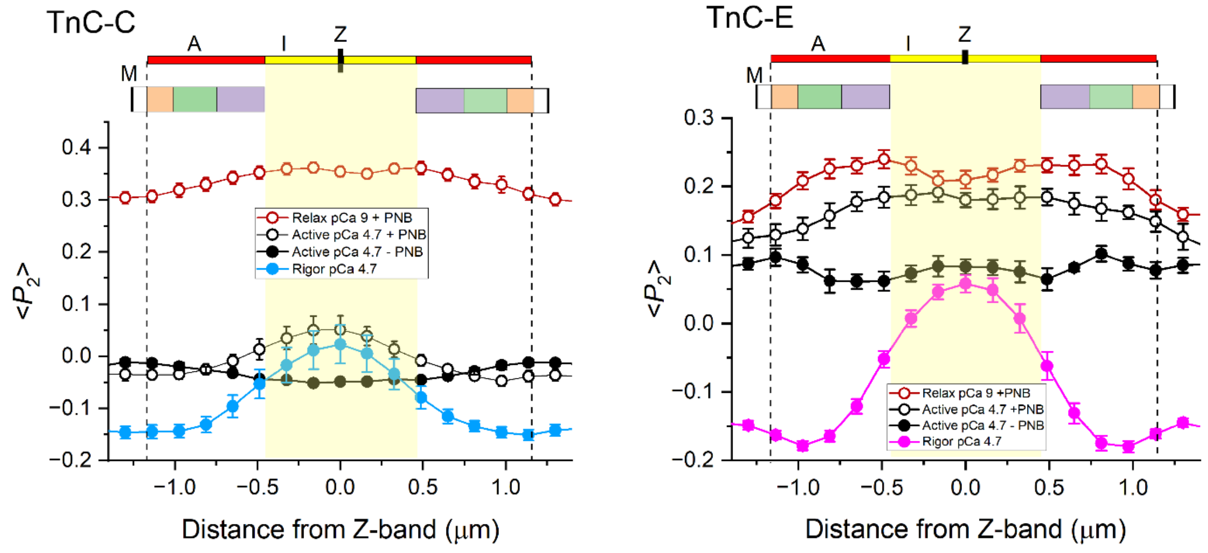


**Fig. S7.** Comparison of the orientation distribution of TnC-C and TnC-E probes along thin filaments in I- (yellow) and A- (red) bands in the sarcomere of relaxed (pCa 9), and active (pCa 4.7) myofibrils in the presence of PNB (data from Fig.5), in active myofibrils (pCa 4.7) in the absence of PNB (data from Fig. 4) and in rigor myofibrils at pCa 4.7 (cyan, data from Fig.1).

**Supplementary Tables**

| | $<P_2>$ | |
|---|---|---|
| **TnC-C** | **FISS (muscle fibres)** | **FPM (myofibrils)** |
| Relax | 0.422 ± 0.021[†] | 0.431 ± 0.012 |
| Rigor pCa 9.0 | 0.207 ± 0.018[†] | 0.232 ± 0.009 |
| Rigor pCa 4.7 | -0.142 ±0.007[†] | -0.088 ± 0.022 |
| Active pCa 4.7 | -0.133 ± 0.004[§] | -0.032 ± 0.005 |

| **TnC-E** | **FISS (muscle fibres)** | **FPM (myofibrils)** |
|---|---|---|
| Relax | 0.196 ± 0.007[†] | 0.210 ± 0.010 |
| Rigor pCa 9.0 | -0.005 ± 0.012[†] | -0.023 ± 0.007 |
| Rigor pCa 4.7 | -0.123 ± 0.014[†] | -0.089 ± 0.010 |
| Active pCa 4.7 | -0.017 ± 0.009[§] | 0.078 ± 0.010 |

| **RLC-E** | **FISS (muscle fibres)** | **FPM (myofibrils)** |
|---|---|---|
| Relax | 0.113± 0.033* | 0.113 ± 0.016 |
| Rigor pCa 9.0 | -0.290 ± 0.024* | -0.220 ± 0.004 |
| Active pCa 4.7 | 0.020 ± 0.012[§] | 0.040 ± 0.011 |

**Table S1.** Comparison of published bulk $<P_2>$ values for TnC and RLC probes measured by wide-field Fluorescence for In-Situ Structure (FISS) in rabbit psoas muscle fibres with corresponding space-averaged $<P_2>$ values measured by Fluorescence Polarisation Microscopy (FPM) in single sarcomeres of skeletal muscle myofibrils from rabbit psoas. Relaxed and rigor data are at 17°C; active data are at 27-30°C.[†]TnC FISS data are from Sun et al. 2006[2]; *RLC FISS data are from Fusi et al. 2015[1]; [§] RLC/TNC FISS data from Brunello et al. 2023[3].

| TnC-C | $<P_2>$ I-band | $<P_2>$ A-band |
|---|---|---|
| Relax | 0.506 ± 0.015 | 0.506 ± 0.015 |
| Rigor pCa 9.0 | 0.415 ± 0.009[a] | 0.171 ± 0.007[a,b] |
| Rigor pCa 4.7 | 0.029 ± 0.050[a,c] | -0.179 ± 0.015[a,b,c] |

| TnC-E | $<P_2>$ I-band | $<P_2>$ A-band |
|---|---|---|
| Relax | 0.193 ± 0.009 | 0.198 ± 0.009 |
| Rigor pCa 9.0 | 0.098 ± 0.011[a] | -0.099 ± 0.006[a,b] |
| Rigor pCa 4.7 | 0.074 ± 0.014[a] | -0.206 ± 0.008[a,b,c] |

| RLC-E | $<P_2>$_P | $<P_2>$_C | $<P_2>$_D |
|---|---|---|---|
| Relax | 0.193 ± 0.019[e] | 0.200 ± 0.023[d] | 0.083 ± 0.020 |
| Rigor pCa 9.0 | -0.194 ± 0.012[a,e,f] | -0.298 ± 0.008[a] | -0.270 ± 0.004[a] |
| Rigor pCa 4.7 | -0.190 ± 0.017[a,e,f] | -0.286 ± 0.006[a] | -0.273 ± 0.005[a] |

**Table S2.** $<P_2>$ values (mean ± SE) for TnC probes in A/I bands and RLC probes in P/C/D zones at 17°C derived from model simulations of the orientation distributions in Fig. 1. Superscripted letters denote significant differences in $<P_2>$ using ANOVA with Tukey's post hoc analysis. [a]P<0.05 when comparing $<P_2>$ in rigor and relaxing conditions. [b]P<0.05 when comparing $<P_2>$ for TnC probes in I- and A-bands. [c]P<0.05 when comparing $<P_2>$ for TnC probes in rigor pCa 9 and pCa 4.7. [d]P<0.05 when comparing $<P_2>$ for RLC-E probe in C- and D-zones. [e]P<0.05 when comparing $<P_2>$ for RLC-E probe in P- and D-zones. [f]P<0.05 when comparing $<P_2>$ for RLC-E probe in P- and C-zones.

| TnC-C | -PNB | | +PNB | |
|---|---|---|---|---|
| | $\langle P_2 \rangle$ I-band | $\langle P_2 \rangle$ A-band | $\langle P_2 \rangle$ I-band | $\langle P_2 \rangle$ A-band |
| pCa 9.0 | 0.478 ± 0.010 | 0.478 ± 0.010 | 0.422 ± 0.010 [a] | 0.398 ± 0.017 [a] |
| pCa 6.6 | 0.355 ± 0.013 | 0.340 ± 0.014 | 0.296 ± 0.016 [a] | 0.268 ± 0.011 [a] |
| pCa 4.7 | -0.061 ± 0.005 | -0.019 ± 0.011 | 0.062 ± 0.035 [a,d] | -0.051 ± 0.012 |

| TnC-E | -PNB | | +PNB | |
|---|---|---|---|---|
| | $\langle P_2 \rangle$ I-band | $\langle P_2 \rangle$ A-band | $\langle P_2 \rangle$ I-band | $\langle P_2 \rangle$ A-band |
| pCa 9.0 | 0.311 ± 0.017 | 0.311 ± 0.017 | 0.255 ± 0.013 | 0.263 ± 0.015 |
| pCa 6.6 | 0.208 ± 0.018 | 0.206 ± 0.017 | 0.236 ± 0.010 | 0.236 ± 0.010 |
| pCa 4.7 | 0.095 ± 0.015 | 0.080 ± 0.010 | 0.208 ± 0.019 [a] | 0.200 ± 0.017 [a] |

| RLC-E | -PNB | | | +PNB | | |
|---|---|---|---|---|---|---|
| | $\langle P_2 \rangle$_P | $\langle P_2 \rangle$_C | $\langle P_2 \rangle$_D | $\langle P_2 \rangle$_P | $\langle P_2 \rangle$_C | $\langle P_2 \rangle$_D |
| pCa 9.0 | 0.222 ± 0.004 | 0.288 ± 0.017 [b,c] | 0.182 ± 0.012 | 0.203 ± 0.011 | 0.273 ± 0.011 [b,c] | 0.203 ± 0.011 |
| pCa 6.6 | 0.154 ± 0.017 | 0.178± 0.022 [b] | 0.106 ± 0.021 | - | - | - |
| pCa 4.7 | 0.046 ± 0.012 | 0.049 ± 0.011 | 0.039 ± 0.011 | 0.185 ± 0.010 [a] | 0.280 ± 0.004 [a,b,c] | 0.175 ± 0.005 [a] |

**Table S3.** $\langle P_2 \rangle$ values (mean ± SE) for TnC probes in A/I bands and RLC probes in P/C/D zones at 30°C derived from model simulations of the orientation distributions in the absence (Fig. 4) and in the presence of 20µM PNB (Fig. 5). Superscripted letters denote significant differences in $\langle P_2 \rangle$ using ANOVA with Tukey's post hoc analysis. [a]P<0.05 when comparing $\langle P_2 \rangle$ in a filament domain in the absence and presence of PNB. [b]P<0.05 when comparing $\langle P_2 \rangle$ for RLC-E probe in C- and D-zones. [c]P<0.05 when comparing $\langle P_2 \rangle$ for RLC-E probe in C- and P-zones. [d]P<0.05 when comparing $\langle P_2 \rangle$ for TnC probes in I- and A-band.

**Supplementary Text**

**Simulation of the spatial distributions of RLC and TnC orientations along thick and thin filaments, respectively, in skeletal myofibrils.**

To calculate the spatial distribution of orientation of RLC and TnC probes along thick and thin filaments, respectively, we assumed a Gaussian distribution for the fluorescence intensity of single fluorophores along the filament:

$$I(x) = e^{\frac{-(x-x_0)^2}{2\sigma^2}}$$

in which $\sigma$ is determined by the point spread function of our optical system and $x_0$ is the position of each probe along the filament. Using fluorescent beads with diameter=170nm as point sources (P7220, Invitrogen) we determined $\sigma$=200nm (FWHM=470nm) for the point spread function of our microscope (Fig. S2), which is consistent with the theoretical value for the spatial resolution (SR) of about 430nm with NA=0.7 and wavelength of fluorescence (λ=620nm) (SR=λ/2NA) .

To calculate the distribution of $<P_2>$ values across the Z-band and the M-band for TnC and RLC probes, respectively, we assumed a homogenous labelling of troponin and myosin along thin and thick filaments, respectively. Specifically, each thin filament contained 30 TnC fluorophores, beginning 70 nm from the centre of the Z-band with a spacing of 38 nm, while each half thick filament contained 49 RLC fluorophores, starting 80 nm from the M-line with a spacing of 14.5 nm (Fig. S2). To account for contributions from adjacent sarcomeres, additional TnC and RLC arrays were placed at distances of ±SL, where SL is the sarcomere length measured experimentally in the myofibril.

The spatial distribution of $<P_2>$ for the RLC probe in each thick filament was calculated as:

$$P_{2\_RLC}(x) = \frac{\sum_0^{11}(< P_2 >_P \cdot I_i ) + \sum_{12}^{29}(< P_2 >_C \cdot I_i ) + \sum_{30}^{48}(< P_2 >_D \cdot I_i)}{\sum_0^{48} I_i + I_{iso}}$$

in which $I_i$ is the fluorescence intensity distribution for the RLC probe at each myosin layer (i), $I_{iso}$ is a constant fluorescence intensity distribution along the sarcomere produced by a population of isotropic ($<P_2>$=0) and non-specifically bound probes, $<P_2>_P$, $<P_2>_C$ and $<P_2>_D$ are the $<P_2>$ values for probes in the P (myosin layers 0-11), C (layers 12-29) and D (layers 30-48) zones of the filament (Fig. S2). $I_{iso}$ was fixed to ~10% of the total maximal fluorescence intensity produced by specifically bound probes. Therefore $<P_2>_P$, $<P_2>_C$ and $<P_2>_D$ are the only free parameters that were adjusted to fit the experimental spatial distributions of RLC orientations.

Similarly, the orientation distributions of $<P_2>$ for the TnC probes in rigor and relaxed myofibrils shown in Fig.1 were calculated as:

$$P_{2\_TnC}(x) = \frac{\sum_0^{11}(< P_2 >_I \cdot I_i) + \sum_{12}^{29}(< P_2 >_A \cdot I_i )}{\sum_0^{29} I_i + I_{iso}}$$

in which $\langle P_2 \rangle_I$, $\langle P_2 \rangle_A$ are $\langle P_2 \rangle$ values for TnC probes in the I- (troponin 0-11) and A-bands (troponin 12-29) of the sarcomere (Fig. S2).
